# Δ40p53 plays a distinct regulatory role in maintaining cellular homeostasis by controlling SGSH expression via miR-4651-5p

**DOI:** 10.1101/2023.04.04.535506

**Authors:** Apala Pal, Pritam Kumar Ghosh, Sahana Ghosh, Sachin Kumar Tripathi, Subrata Patra, Debjit Khan, Arindam Maitra, Saumitra Das

**Author notes:** Address for correspondence: Dr. Saumitra Das, Professor, Department of Microbiology and Cell Biology, Indian Institute of Science, Bangalore-560012, India.

## Abstract

Δ40p53, the only translational isoform of p53, modulates the full-length p53 (FLp53) activity and independently regulates targets such as the miR-186-5p–YY1 axis. To identify additional miRNAs regulated by Δ40p53, we performed small RNA sequencing. We found that overexpression of Δ40p53, but not FLp53, significantly downregulated miR-4671-5p. Expression of both isoforms at varying ratios revealed that miR-4671-5p may be modulated by FLp53 in a Δ40p53-dependent manner. *In silico* analysis identified SGSH (N-sulfoglucosamine sulfohydrolase) as a potential miR-4671-5p target. SGSH expression showed inverse correlation with miR-4671-5p in cancer datasets and prognostic significance. SGSH mRNA and protein levels were reduced upon miR-4671-5p overexpression or siΔ40p53 treatment, confirming regulatory linkage. Functionally, miR-4671-5p overexpression induced intra-S-phase cell cycle arrest, implicating SGSH in cell cycle regulation. These results reveal a novel Δ40p53–miR-4671-5p–SGSH axis that impacts cell cycle progression and may contribute to cancer outcomes. Our findings highlight the distinct regulatory role of Δ40p53, independent of FLp53, in maintaining cellular and metabolic homeostasis via miRNA-mediated mechanisms.

## INTRODUCTION

Genome integrity relies on rapid recognition and repair of DNA damage. Central to this response is p53, which halts the cell cycle, limits proliferation of damaged cells, and is frequently inactivated by mutation in cancer (Kops et al., 2005; Rivlin et al., 2011). p53 activity is further shaped by 12 naturally occurring isoforms that differ at the N- and C-termini (Bourdon, 2007; Khoury & Bourdon, 2011). Their relative abundance varies across normal tissues and tumours, altering transcriptional output and cellular fate (Bourdon et al., 2005; Vieler & Sanyal, 2018). Δ40p53 is the sole translational isoform, produced from an internal ribosome-entry site (IRES) within the TP53 5′-UTR (Ray et al., 2006). A second IRES yields full-length p53 (FLp53); the two proteins share oligomerisation domains and form homo- and hetero-tetramers (Ghosh et al., 2004). IRES usage is stress- and cell-cycle-phase dependent: translation of FLp53 peaks at the G2/M transition, whereas Δ40p53 is preferentially produced at G1/S (Ray et al., 2006; Khan et al., 2015).

Functionally, Δ40p53 fine-tunes FLp53 activity yet also acts independently. It retains the second transactivation domain of p53, driving gene expression in p53-null cells (Maier et al., 2004; Ohki et al., 2007). Reported activities include induction of 14-3-3σ and G2 arrest (Bourougaa et al., 2010), activation of pro-apoptotic BAX and GADD45 (Steffens Reinhardt et al., 2020), modulation of the Nanog–IGF1 axis during stem-cell differentiation (Ungewitter & Scrable, 2010), and control of β-cell proliferation and glucose homeostasis (Hinault et al., 2011). At baseline, Δ40p53 and FLp53 suppress migration and proliferation in breast cancer (Zhang et al., 2022). Conversely, Δ40p53 can promote survival by transactivating netrin-1 (Sun et al., 2021). Growing evidence links Δ40p53 to non-coding RNA networks. We previously showed that Δ40p53—but not FLp53— up-regulates miR-186-5p, which represses the oncogenic transcription factor YY1 and limits proliferation (Katoch et al., 2021). Δ40p53 also controls the lncRNA LINC00176, influencing multiple miRNAs and mRNAs that govern cell fate (Pal et al., 2023). These findings suggest that isoform-specific miRNA programmes are key to Δ40p53 function, yet a systematic catalogue of such miRNAs is lacking.

We performed small-RNA sequencing to globally identify miRNAs differentially regulated by Δ40p53 and FLp53. Using lung (H1299) and colon (HCT116) cancer cell lines engineered to express FLp53, Δ40p53, or both, we discovered miR-4671-5p as a Δ40p53-specific target. Down-stream analyses revealed its repression of N-sulfoglucosamine sulfohydrolase (SGSH), linkage to S-phase arrest, and prognostic significance in multiple tumours. Here, we delineate the Δ40p53–miR-4671-5p–SGSH axis and its impact on cell-cycle control, providing fresh insight into isoform-specific p53 signalling.

## MATERIALS AND METHODS

### Cell lines and transfections

Three human cell lines were used: H1299 (lung adenocarcinoma, p53-null), HCT116 p53+/+ (referred to as HCT116+/+), and HCT116 p53–/– (referred to as HCT116–/–), which endogenously expresses only Δ40p53. Cells were cultured in Dulbecco’s Modified Eagle Medium (DMEM, Sigma) supplemented with 10% fetal bovine serum (FBS, GIBCO, Invitrogen) and 1% penicillin/streptomycin. Transfections were performed using Lipofectamine 2000 and Turbofectamine (Invitrogen) in Opti-MEM (GIBCO) according to the manufacturer’s instructions. After 4 hours, transfection medium was replaced with complete DMEM, and cells were harvested at indicated time points.

Plasmid constructs included pGFP-hp-p53-5′UTR vectors expressing FLp53, Δ40p53, or both (14A). These were generously provided by Dr. Robin Fahraeus (INSERM, France). To overexpress miR-4671-5p, the mature sequence was cloned into the pSUPER vector. For knockdown studies, a 30 nM siRNA targeting the 3′UTR of TP53 (IDT) was used, which downregulates both FLp53 and Δ40p53; a non-targeting control siRNA (Dharmacon) was used in parallel.

### Small RNA Sequencing

#### RNA extraction

Total RNA was isolated from H1299 cells transfected with vector, FL p53, Δ40p53, or 14A using TRIzol followed by the PureLink kit (Ambion). RNA quality (Bioanalyzer 2100, Agilent) and quantity (NanoDrop, Qubit) were assessed; only samples with RIN > 7 were used for library preparation.

#### Library preparation and sequencing

One µg of high-quality RNA was processed with the Illumina TruSeq Small RNA kit. 3′/5′ adapters were ligated, cDNA was synthesised, PCR-amplified, and fragments of 145–160 bp were gel-purified. Libraries were pooled and sequenced (1 × 50 bp) on an Illumina HiSeq-2500.

#### RNA-seq data analysis

FASTQ files were trimmed with Cutadapt v1.8.1 and mapped to hg19 using miRDeep2. Known and novel miRNAs were quantified as reads-per-million (RPM); multi-precursor miRNAs were averaged. Differential expression was determined with DESeq2.

### Western Blot analysis

Cells were lysed in RIPA buffer, and protein concentrations were determined via the Bradford assay. Equal amounts of lysate were resolved by SDS-PAGE and transferred to nitrocellulose membranes. Primary antibodies used were: anti-p53 (CM1; kindly provided by Dr. Robin Fahraeus (INSERM, France) and Prof. J.C. Bourdon (University of Dundee, UK), anti-SGSH (Abclonal A8148), anti-CDK11B (Ablconal A12830), and anti-CDK5R1 (Ablconal A14497). Blots were developed using HRP-conjugated secondary antibodies (Sigma) and detected with enhanced chemiluminescence (ECL).

### RNA isolation and Real-time PCR

Total RNA was extracted using TRI Reagent™ (Sigma) and treated with DNase I (Promega) to remove DNA contamination. RNA was purified by acidic phenol–chloroform extraction and ethanol precipitation, then quantified using a Nano-spectrophotometer. cDNA was synthesized from 2–5 µg total RNA for mRNAs or 50 ng for miRNAs using gene-specific primers and Revertaid™ MMLV RT (Thermo Scientific) at 42 °C for 1 hour.

### Cell cycle Analysis

HCT116–/– cells were transfected with miR-4671-5p or miR-34a-5p overexpression constructs. After 48 h, cells were fixed in methanol, treated with RNase A (10 µg/mL), stained with propidium iodide (1 µg/mL), and analyzed on a BD FACSCalibur flow cytometer.

### Bioinformatics and statistics

dbDEMC2.0 has been used to perform a meta-profiling of selected miRNAs (Xu et al., 2022). TargetScan 7.2 has been used to predict target mRNAs of miRNAs (Agarwal et al., 2015). miRDB has been used to predict target miRNAs (Chen and Wang, 2020; Liu and Wang, 2019). GO-term analysis was done on PANTHER release 17.0. Kaplan-Meier survival analysis was done on R2: Genomics Analysis and Visualization Platform. Heat map for Figure 1E has been generated using Heatmapper. Data were analyzed using two-tailed Student’s t-test; p≤0.05 (*) or p≤0.01 (**) or p≤ 0.001(***)

## RESULTS

### Small RNA sequencing reveals miRNA targets of p53 and Δ40p53

H1299 cells, which lack endogenous p53 isoforms, were transfected to express FLp53, Δ40p53, both isoforms (14A), or GFP (control) **(Figure 1A–C)**. Total RNA was extracted, quality-checked, and subjected to small RNA sequencing. From the dataset, 135 miRNAs with consistent unidirectional fold changes across replicates were selected. A heatmap visualized their expression profiles under each isoform condition **(Figure 1D)**. A statistically significant cluster of differentially expressed miRNAs (p ≤ 0.05) was identified **(Figure 1E)**. Among these, four miRNAs stood out for their isoform-specific regulation. miR-4671-5p was significantly downregulated by Δ40p53 but unaffected by FLp53. In contrast, miR-301b-5p was reduced only in FLp53-expressing cells. miR-34a-5p, a known p53 target (Zhang et al., 2019) was upregulated in all isoform conditions and served as a positive control. Similarly, miR-548ae-5p was upregulated across FLp53, Δ40p53, and 14A conditions, indicating potential co-regulation. These results reveal distinct and overlapping miRNA regulatory profiles for FLp53 and Δ40p53, with miR-4671-5p emerging as a prominent Δ40p53-specific target for further investigation.

**Figure 1.**
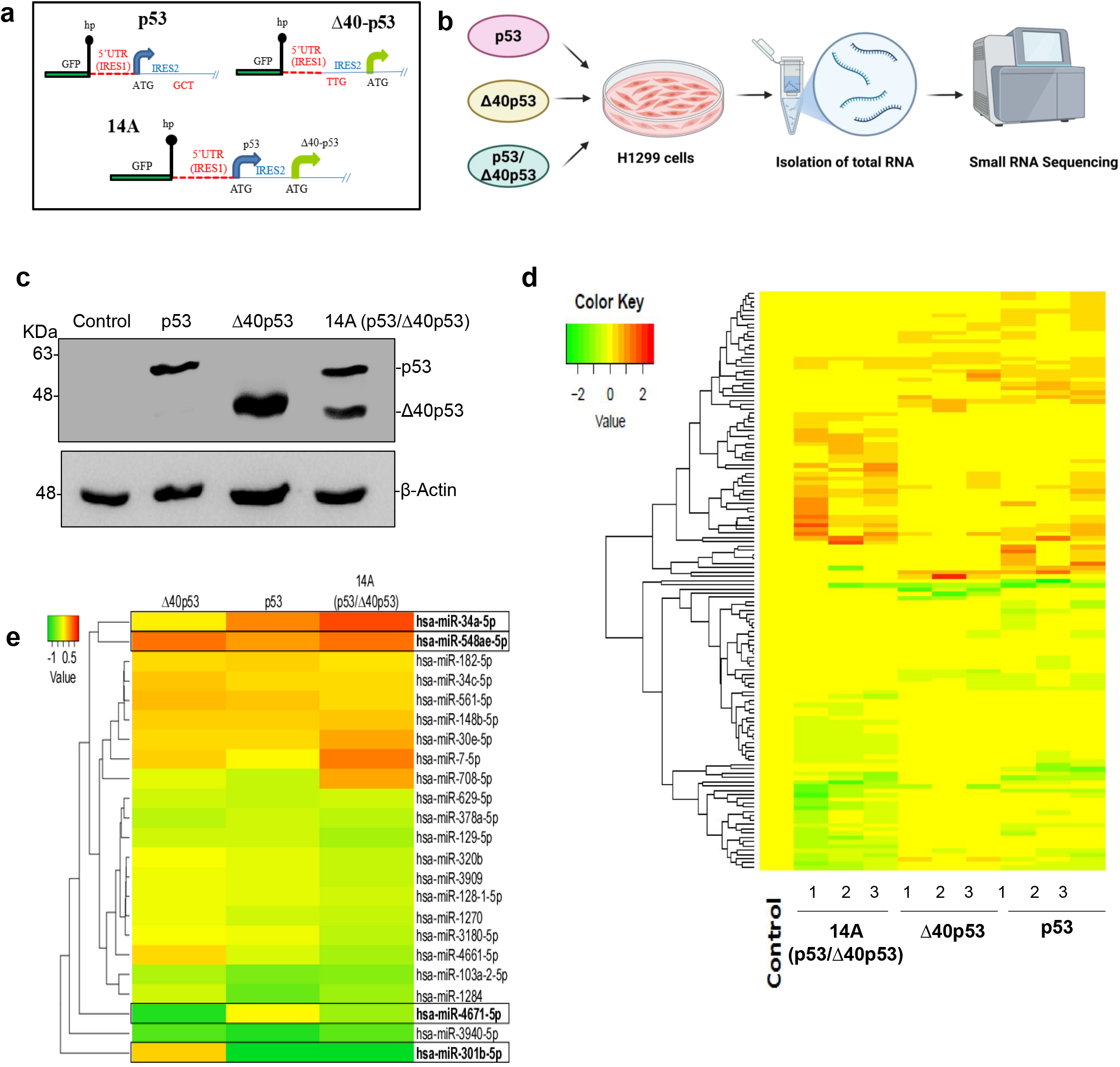
Regulation of miRNA expression by p53 and its isoform Δ40p53. (A) Schematic of constructs used in study (B) Experimental setup prior to RNA sequencing. (C) Western blot analysis of cell extracts from H1299 cells expressing control, p53 only, Δ40p53 only and 14A construct, probed with CM1 after 48 h. (D) The cluster of differentially expressed miRNAs obtained from the small RNA sequencing data set. (E) The compressed cluster of significant differentially expressed miRNAs with p-value ≤ 0.05 (generated using Heatmapper).

### Selection and validation of miRNAs for further studies

To validate small RNA sequencing results, H1299 cells were transfected with FLp53, Δ40p53, 14A (both isoforms), or control vector (**Figure 2A**), and miRNA levels were measured by qRT-PCR. miR-4671-5p was significantly downregulated by Δ40p53 and 14A, but not by FLp53 alone (**Figure 2B**). In contrast, miR-34a-5p and miR-548ae-5p were consistently upregulated across all isoform conditions (**Figure 2C–D**). miR-301b-5p showed mild induction by Δ40p53 and 14A, but not FLp53 (**Figure 2E**). Expression patterns for miR-4671-5p, miR-548ae-5p, and miR-34a-5p matched the sequencing data (**Figure 2F**); miR-301b-5p was excluded from further analysis.

**Figure 2.**
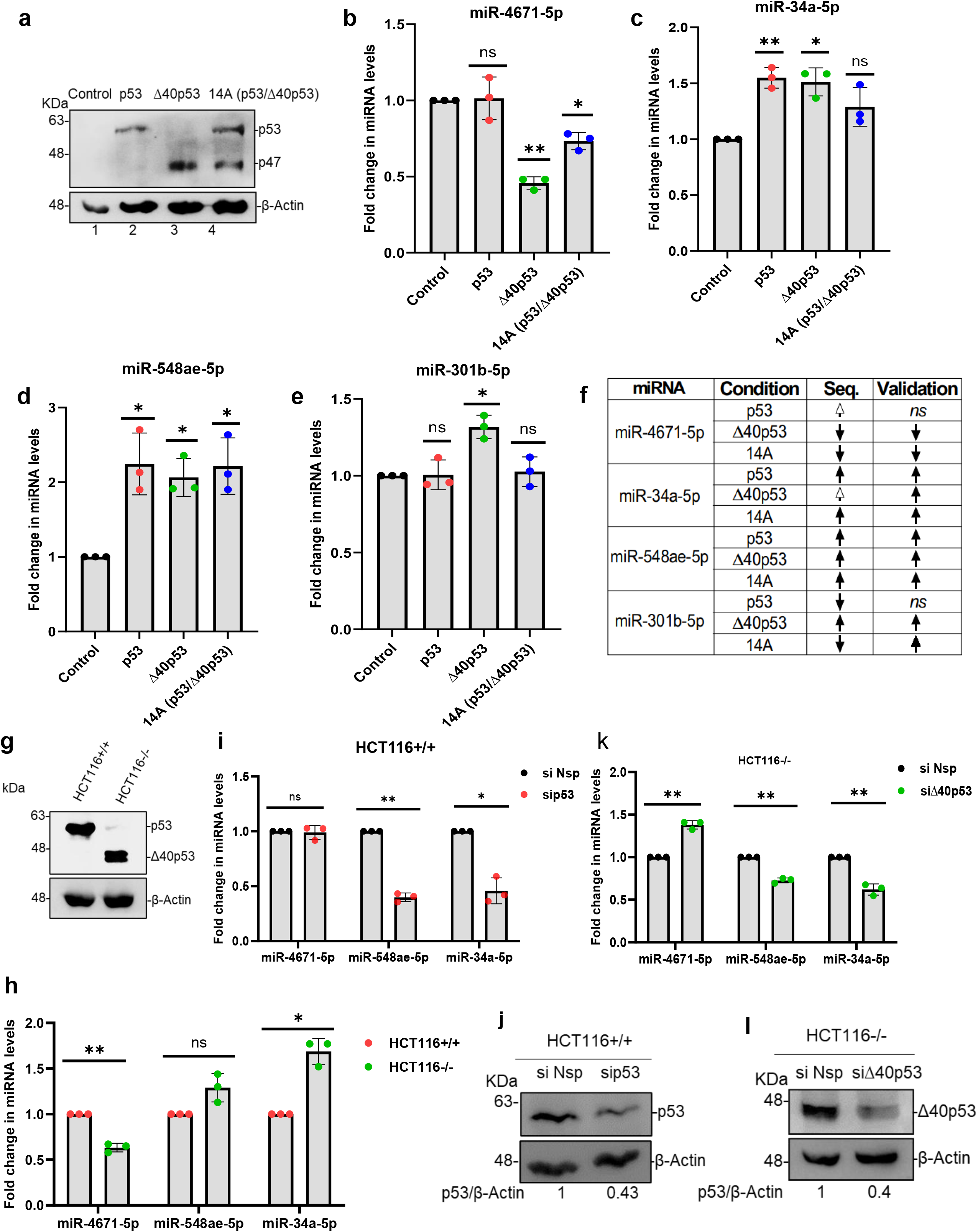
Selection and validation of miRNAs for further studies. (A) Western blot analysis of cell extracts from H1299 cells expressing control, p53 only, Δ40p53 only and 14A construct, probed with CM1 after 48 h. (B-E) Quantitative PCR for validation of miR-4671-5p, miR-34a-5p, miR-548ae-5p, miR-301b-5p, respectively, in H1299 cells expressing control, p53 only, Δ40p53 only and 14A construct. (F) Table for comparative analysis of fold changes obtained in sequencing versus validation result. (G) Western blot analysis of cell extracts from HCT116+/+ and HCT116-/-cells probed with CM1. (H) Quantitative PCR for validations of miR-4671-5p, miR-34a-5p and miR-548ae-5p in HCT116+/+ and HCT116-/-cells. (I) Quantitative PCR of miR-4671-5p, miR-34a-5p and miR-548ae-5p in HCT116+/+ cells transfected with si p53 (30nM) and non-specific si (si Nsp). (J) Western blot analysis of cell extracts from HCT116+/+ cells transfected with either si p53 (30nM) and non-specific si (si Nsp), probed with CM1. (K) Quantitative PCR of miR-4671-5p, miR-34a-5p and miR-548ae-5p in HCT116-/-cells transfected with si Δ40p53 (30nM) and non-specific si (si Nsp). (L) Western blot analysis of cell extracts from HCT116-/-cells transfected with siΔ40p53 (30nM) and non-specific si (si Nsp), probed with CM1 The criterion for statistical significance was *p≤*0.05 (*) or *p≤*0.01 (**) or p*≤* 0.001(***)

To assess physiological relevance, we examined these miRNAs in colon cancer cell lines HCT116+/+ (predominantly FLp53) and HCT116–/– (only Δ40p53) (**Figure 2G**). At baseline, HCT116–/– cells showed lower miR-4671-5p and higher miR-548ae-5p and miR-34a-5p levels than HCT116+/+ cells (**Figure 2H**). siRNA-mediated knockdown of p53 in HCT116+/+ cells reduced miR-548ae-5p and miR-34a-5p but did not significantly affect miR-4671-5p (**Figure 2I–J**). Conversely, Δ40p53 knockdown in HCT116–/– cells increased miR-4671-5p and decreased miR-548ae-5p and miR-34a-5p (**Figure 2K–L**). These results suggest that Δ40p53 specifically suppresses miR-4671-5p and enhances miR-548ae-5p and miR-34a-5p expression, while FLp53 does not regulate miR-4671-5p.

Given that p53 isoform expression varies widely across cancer types (Vieler and Sanyal, 2018), we next examined whether the expression patterns of these miRNAs—particularly miR-4671-5p—also show variability in clinical samples. To explore this, we analyzed publicly available cancer transcriptome datasets.

### Regulation of miRNAs by Δ40p53 and p53

Bioinformatic analysis using dbDEMC 2.0 revealed that miR-34a-5p is widely expressed in many cancers, while miR-4671-5p expression is restricted to colon, lung, and pancreatic cancers. In contrast, miR-548ae-5p was undetectable in this dataset (**Figure 3A**). These observations suggest that the differential expression of these miRNAs may reflect the tissue-specific abundance of p53 and Δ40p53. To test this, we overexpressed p53 and Δ40p53 in varying ratios in H1299 cells and analyzed miRNA expression (**Figure 3B**). Expression of miR-34a-5p was induced by either isoform alone; however, certain combinations of both isoforms reduced its expression, suggesting dose-dependent co-regulation (**Figure 3C**), consistent with previous reports of p53/Δ40p53 interactions (Ghosh et al., 2004).

**Figure 3.**
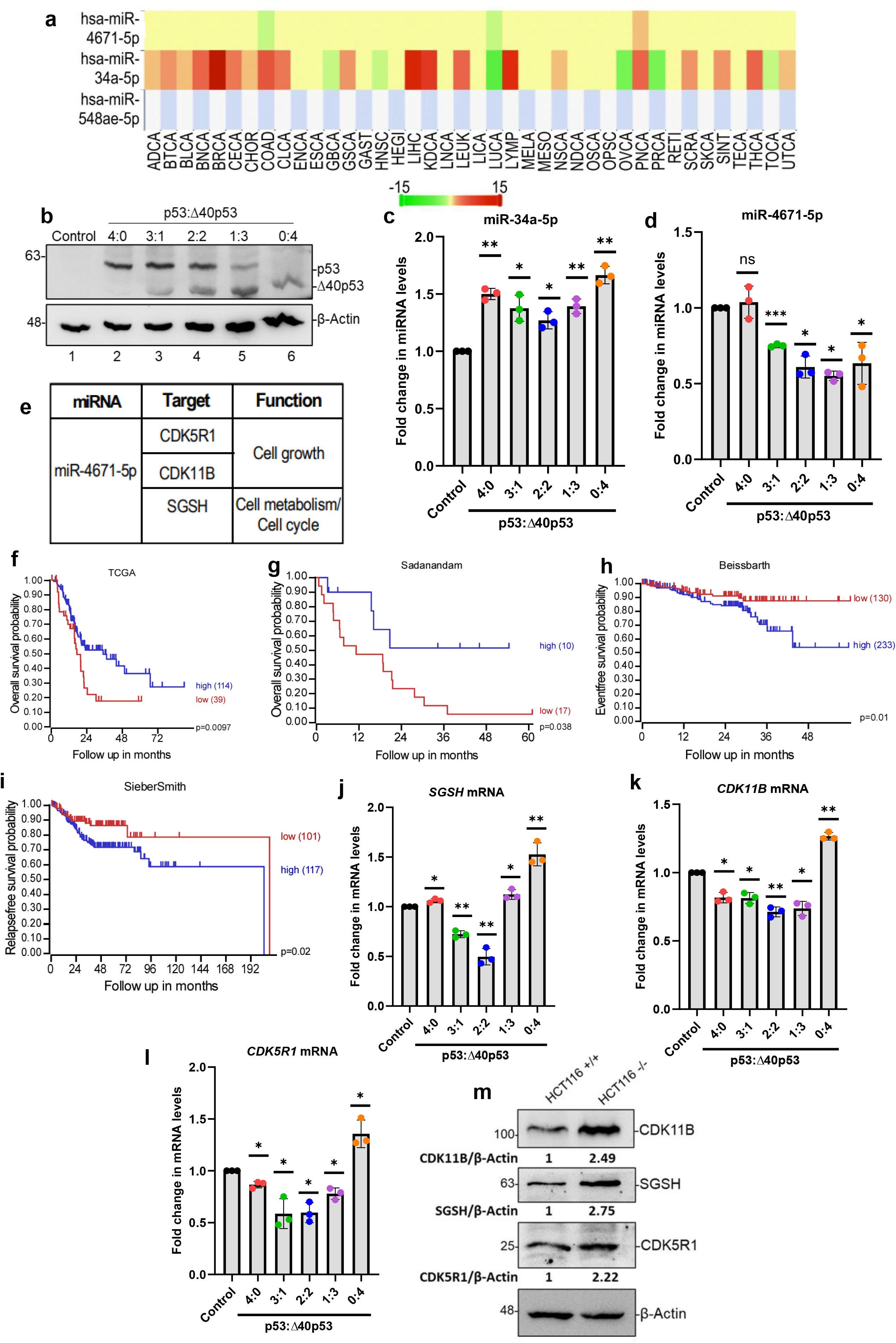
Regulation of miRNAs by Δ40p53 and p53. (A) Differential expression Profile of miRNAs in cancer vs. normal obtained from dbDEMC 3.0 database. (B) Western blot analysis of cell extracts from H1299 transfected cells with different ratios of p53 and Δ40p53 probed with CM1. (C-D) Quantitative PCR of miR-34a-5p and miR-4671-5p respectively in H1299 transfected with different ratios of p53 and Δ40p53. (E) Table for miR-4671-5p targets selected in the study. (F-I) Kaplan-Meier estimates of survival for pancreatic adenocarcinoma (F-G) and colon tumor (H-I) classified by SGSH mRNA expression levels. The number of patients at risk is indicated for time increments of 12 or 24 months. p values were calculated using a log rank test. (J-L) Quantitative PCR of mRNA targets (SGSH, CDK11B and CDK5R1) in H1299 transfected cells with different ratios of p53 and Δ40p53. (M) Western blot analysis of cell extracts from HCT116+/+ and HCT116-/-cells probed with CM1, SGSH, CDK11B, CDK5R1 antibodies. The criterion for statistical significance was *p≤*0.05 (*) or *p≤*0.01 (**) or p*≤* 0.001(***).

Notably, FLp53 overexpression alone did not significantly affect miR-4671-5p levels, whereas increasing Δ40p53 expression—either alone or in combination—consistently suppressed it (**Figure 3D**). The 14A construct mimicked this effect, indicating that p53 may modulate Δ40p53-mediated regulation of miR-4671-5p but is not itself sufficient to alter its expression. These data reinforce that Δ40p53 is the primary regulator of miR-4671-5p and that isoform balance is critical for miRNA output.

### Cellular role of miR-4671-5p

Because miR-4671-5p was uniquely repressed by Δ40p53, we explored its downstream functions. TargetScan 7.2 and miRDB yielded 786 non-redundant putative targets, with GO enrichment for ‘protein-serine-kinase activity’ indicating that many predicted targets of miR-4671-5p may be involved in cell cycle regulation. From this list we prioritised two cell-cycle kinases, CDK11B and CDK5R1 for experimental follow-up (**Figure 3E; Figure S1**). Additionally, we prioritized SGSH (N-sulfoglucosamine sulfohydrolase), the top-scoring miR-4671-5p target in TargetScan, despite its uncharacterized role in cancer.

SGSH is a lysosomal enzyme whose deficiency causes mucopolysaccharidosis IIIA (MPS IIIA), leading to GAG and heparan sulfate accumulation and resulting in severe neurological and skeletal pathology (Li et al., 2018). Although SGSH has recently been linked to monkeypox and interstitial lung disease (Anuraga et al., 2024; Ye et al., 2022), its role in tumor biology remains unknown. Analysis of patient datasets revealed that low SGSH expression correlates with poor survival in pancreatic cancer, while high SGSH is associated with poor outcomes in colon tumors (**Figure 3F–I; Fig. S1, Table 3**). These trends are opposite to miR-4671-5p levels observed in the same cancers (**Figure 3A**), suggesting a biologically relevant interaction.

We next validated the regulation of these targets under varying p53/Δ40p53 expression in H1299 cells. *SGSH, CDK11B*, and *CDK5R1* mRNA levels were upregulated only under Δ40p53 overexpression (**Figure 3J–L**). Similarly, in HCT116–/– cells (expressing endogenous Δ40p53), these proteins were more abundant than in HCT116+/+ cells (**Figure 3M**), consistent with lower miR-4671-5p expression (**Figure 3A**). Partial knockdown of Δ40p53 reduced expression of all three targets (**Figure 4A–B**), while overexpression of miR-4671-5p in HCT116–/– cells led to reduced SGSH, CDK11B, and CDK5R1 mRNA levels, most prominently for SGSH (**Figure 4C–D**). Corresponding decreases in SGSH protein were observed under both conditions (**Figure 4E–F**).

**Figure 4.**
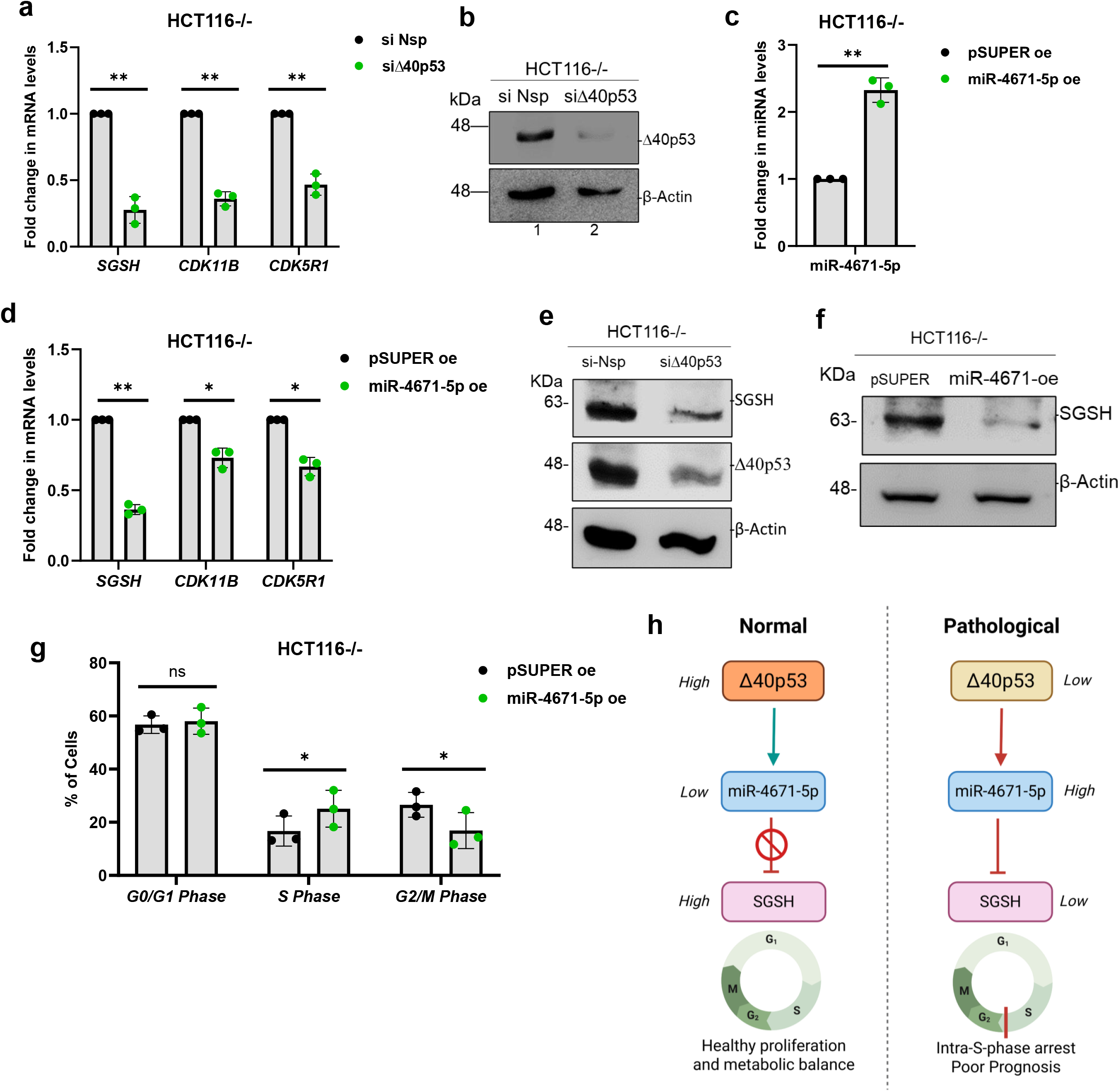
role of miR-4671-5p. (A) Quantitative PCR of SGSH, CDK11B and CDK5R1 in HCT116-/-cells transfected with si Δ40p53 (30nM) and non-specific si (si Nsp). (B) Western blot analysis of cell extracts from HCT116-/-cells transfected with siΔ40p53 (30nM) and non-specific si (si Nsp), probed with CM1. (C) Quantitative PCR of miR-4671-5p in HCT116-/-cells transfected with miR-4671-5p overexpression construct. (D) Quantitative PCR of SGSH, CDK11B and CDK5R1 in HCT116-/-cells transfected with miR-4671-5p overexpression construct. (E) Western blot analysis of cell extracts from HCT116-/-cells transfected with siΔ40p53 (30nM) and non-specific si (si Nsp), probed with CM1 and SGSH. (F) Western blot analysis of cell extracts from HCT116-/-cells transfected with miR-4671-5p overexpression construct, probed with SGSH. (G) Analysis of different cell cycle phases of fixed cell extracts from HCT116-/-cells transfected with miR-4671-5p overexpression construct. The criterion for statistical significance was *p≤*0.05 (*) or *p≤*0.01 (**) or p*≤* 0.001(***). (H) Left (Normal condition): Under physiological conditions, high levels of the p53 translational isoform Δ40p53 suppress the expression of miR-4671-5p. Reduced levels of miR-4671-5p relieve its inhibitory effect on SGSH, leading to elevated SGSH expression. High SGSH levels support normal cell cycle progression through G1, S, G2, and M phases. Right (Pathological condition): In conditions where Δ40p53 expression is low, miR-4671-5p is upregulated. Increased miR-4671-5p inhibits SGSH expression, resulting in reduced SGSH levels. This suppression contributes to intra-S-phase cell cycle arrest, which is associated with impaired proliferation control and poor cancer prognosis.

Finally, miR-4671-5p overexpression caused significant S-phase accumulation and reduced G2-phase cells, indicating intra-S phase arrest (Fig. 4G). Together, these findings establish that Δ40p53 suppresses miR-4671-5p to maintain SGSH and other cell cycle regulators, enabling proper S-phase progression.

## DISCUSSION

MicroRNAs (miRNAs) play critical roles in regulating development, cell cycle progression, and survival. Their dysregulation is a common feature in cancer (Israel et al., 2009). While many miRNAs are known to be transcriptionally regulated by full-length p53 (FLp53) (Takwi and Li, 2009), the contribution of p53 isoforms—especially Δ40p53—to miRNA regulation has only recently been explored. We previously demonstrated that Δ40p53 selectively upregulates miR-186-5p, which targets and represses the oncogene YY1, leading to reduced cell proliferation (Katoch et al., 2021). Despite this, the full functional scope of Δ40p53 remains understudied, even though it has been implicated in key cellular processes such as apoptosis, senescence, migration, and the cell cycle (Steffens Reinhardt et al., 2020). A key open question is: which non-coding RNAs are specifically regulated by Δ40p53, and how might these contribute to its distinct roles? This is particularly relevant in cancers, where differential isoform expression (Vieler and Sanyal, 2018) may reshape downstream gene regulatory networks.

Here, we identified several miRNAs regulated by Δ40p53, either uniquely or in parallel with p53. Among these, miR-4671-5p was exclusively repressed by Δ40p53, prompting further investigation. Overexpression of Δ40p53 significantly reduced miR-4671-5p levels, whereas p53 alone had no effect. Interestingly, the dual-isoform construct (14A) also reduced miR-4671-5p expression (Figure 2B), suggesting that p53 may enhance Δ40p53-mediated repression. Similarly, varying p53/Δ40p53 ratios consistently decreased miR-4671-5p expression in the presence of both isoforms (Figure 3D). These findings indicate that relative isoform abundance modulates miR-4671-5p levels, which could contribute to context-dependent regulation of downstream targets in different cancers.

To explore the downstream role of miR-4671-5p, we initially selected SGSH, CDK11B, and CDK5R1 as candidate targets based on bioinformatic predictions and GO enrichment analysis (**Figure S1, Table 2**). Among these, SGSH was prioritized for further investigation due to its strong prediction score, metabolic function, and uncharacterized role in cancer. SGSH expression also showed prognostic significance: low SGSH associated with poor survival in pancreatic cancer, whereas high SGSH correlated with worse outcome in colon cancer, glioblastoma (**Figure 3F-I; Figure S1, Table 3**) and other datasets. Notably, SGSH and miR-4671-5p levels were inversely related across these tumours (**Figure 3A**), supporting functional linkage. The bidirectional prognostic pattern suggests SGSH is neither a classic oncogene nor tumour suppressor but may operate within tumour-specific rewiring of the Δ40p53–miR-4671-5p–SGSH axis—an area that merits deeper investigation.

**Table 1.**

| Database | Target Scan database | miRDB |
| --- | --- | --- |
| miRNA | Number of targets | Number of targets |
| miR-4671-5p | 780 | 46 |

**Table 2.**

| “GO molecular function complete” terms | *Homo sapiens* (REF) # genes | hsa-miR-4671-5p target list # genes | Expected # genes | Fold Enrichment | +/- | raw P value | FDR |
| --- | --- | --- | --- | --- | --- | --- | --- |
| protein serine kinase activity | 364 | 32 | 13.67 | 2.34 | + | 2.77E-05 | 2.30E-02 |
| protein serine/threonine kinase activity | 433 | 35 | 16.26 | 2.15 | + | 7.52E-05 | 4.16E-02 |
| molecular adaptor activity | 488 | 38 | 18.32 | 2.07 | + | 7.07E-05 | 4.40E-02 |
| ion binding | 6088 | 283 | 228.57 | 1.24 | + | 3.63E-05 | 2.58E-02 |
| binding | 16594 | 694 | 623.01 | 1.11 | + | 1.01E-11 | 5.01E-08 |
| protein binding | 14448 | 618 | 542.44 | 1.14 | + | 1.83E-09 | 4.56E-06 |
| Unclassified | 2297 | 44 | 86.24 | 0.51 | - | 2.92E-07 | 3.64E-04 |
| olfactory receptor activity | 441 | 2 | 16.56 | 0.12 | - | 2.54E-05 | 2.53E-02 |

**Table 3.**

| Cancer/tumor | Dataset | SGSH level (is poor) | p values |  |  |  |  |
| --- | --- | --- | --- | --- | --- | --- | --- |
|  |  |  | Scan (low/high) | Median (Low/high) | Q1 v Rest (Low/high) | Rest v Q4 (Low/high) | Q1 v Q4 (Low/high) |
| Mixed Pancreatic Adenocarcinoma | tcga | LOW | 2.8e-03 (38/115) |  | 9.7 e-03 (39/114) |  |  |
| Mixed Pancreatic Adenocarcinoma | tcga | LOW | 3.1e-05 (8/138) |  |  |  |  |
| Mixed Tumor Pancreas (Exon) | Zhang | HIGH | 0.035 (73/17) |  |  |  |  |
| Mixed Pancreatic | Sadanandam | LOW | 0.038 (17/10) |  |  |  |  |
| Mixed colon adenocarcinoma | TCGA | HIGH | 0.0195 (32/123) |  |  |  |  |
| Mixed colon adenocarcinoma (2022-v32) | TCGA | HIGH | 0.026 (29/437) |  |  |  |  |
| Tumor colon | Sieber | HIGH | 0.023 (181/45) |  |  |  |  |
| Tumor colon | SieberSmith | HIGH | 0.059 (116/170) |  |  |  |  |
| Tumor colon | Smith | HIGH** | 0.011 (98/134) | 0.031 (117/115) |  |  |  |
| Tumor colon | Sveen | HIGH | 3.1e-03 (20/300) |  |  |  |  |
| Tumor colon | Beissbarth | HIGH | 0.01 (130/233) |  |  |  |  |
| Tumor colon | Beauchamp | LOW | 0.012 (53/103) |  |  |  |  |
| Tumor colon | Tsunoda | HIGH | 5.0e-03 (78/11) |  |  |  |  |
| Tumor colon | Marisa | HIGH | 1.1e-03 (539/18) |  |  | 0.061 (418/139) |  |
| Tumor colon | SieberSmith | HIGH | 0.02 (101/177) |  |  |  |  |
| Tumor glioblastoma | TCGA | HIGH | 4.0e-03 (58/106) |  |  |  |  |
| Tumor glioblastoma | TCGA | HIGH | 5.8e-04 (12/365) | 0.021 (189/188) | 0.012 (95/282) |  | 0.051 (95/95) |
| Tumor glioblastoma | TCGA | HIGH | 2.3e-04 (18/486) | 1.6e-03 (253/251) | 3.0e-03 (127/377) |  | 3.8e-03 (127/127) |
| Tumor Neuroblastoma | Kocak | LOW | 7.1e-12 (141/335) | 3.9e-08 (239/237) | 1.6e-08 (120/356) | 1.8 e-04 (357/119) | 6.4e-08 (120/120) |
| Tumor Neuroblastoma | Maris | LOW | 7.6e-03 (68/33) |  |  | 0.022 (76/25) | 0.039 (26/26) |
| Tumor Neuroblastoma | SEQC | LOW | 3.0e-17 (184/314) | 2.3e-11 (249/249) | 5.5e-12 (125/373) | 5.0e-06 (374/124) | 2.3e-11 (125/125) |
| Tumor Neuroblastoma | NRC | LOW | 4.4e-05 (60/223) | 1.1e-03 (142/141) | 5.8e-04 (71/212) | 5.8e-03 (213/70) | 2.4e-04 (71/71) |
| Tumor Neuroblastoma | Asgharzadeh | LOW | 0.016 (111/136) | 0.042 (124/123) |  |  |  |
| Tumor Neuroblastoma | Cangelosi | LOW | 1.9e-16 (303/483) | 1.6e-12 (393/393) | 9.4e-12 (197/589) | 3.6 e-06 (590/196) | 9.0 e-12 (197/197) |
| Tumor Neuroblastoma | Seeger | LOW | 1.1e-03 (25/77) |  | 3.2e-03 (26/76) |  | 0.019 (26/26) |
| Tumor Neuroblastoma | Versteeg | LOW | 0.054 (47/41) |  |  |  |  |
| Tumor Neuroblastoma | Westermann | LOW | 9.1e-05 (68/71) | 4.4e-04 (70/69) |  | 0.015 (105/34) |  |
| Cancer/tumor | Dataset | hsa-miR-4671-5p (is poor) | p values |  |  |  |  |
|  |  |  | Scan (low/high) | Median (Low/high) | Q1 v Rest (Low/high) | Rest v Q4 (Low/high) | Q1 v Q4 (Low/high) |
| Tumor Neuroblastoma | Bell | HIGH | 0.058 (79/16) |  |  |  |  |

Overexpression of Δ40p53 increased the mRNA levels of all three targets—SGSH, CDK11B, and CDK5R1 (**Figure 3J–L**)—with SGSH showing the **most pronounced upregulation**. Protein levels of all three were also higher in cells with endogenous Δ40p53 (**Figure 3M**). Although co-expression of FLp53 and Δ40p53 reduced miR-4671-5p levels (**Figure 3D**), this was not mirrored at the target mRNA level (**Figure 3J–L**), likely due to additional miRNAs regulated by the isoform combination influencing target expression. To validate the Δ40p53– miR-4671-5p–mRNA axis, we overexpressed miR-4671-5p or silenced Δ40p53, both of which led to reduced expression of all three targets, with **SGSH showing the most substantial decrease** (**Figure 4A, D**). This trend was confirmed at the protein level, where SGSH levels dropped consistently under both conditions (**Figure 4E, F**), highlighting SGSH as the **primary and most responsive target** within the Δ40p53–miR-4671-5p regulatory network.

To assess the physiological relevance of SGSH–miR-4671-5p regulation, we considered SGSH’s established role in lysosomal degradation of heparan sulfate (HS), a glycosaminoglycan (Boado et al., 2018; Griffin and Gloster, 2017). In MPS Type IIIA, SGSH mutations impair this process, leading to intracellular accumulation of heparan sulfate. Notably, flow cytometry analyses in MPS models have revealed disrupted cell cycle profiles, including elevated S-phase and reduced G2-phase populations (Brokowska et al., 2022), suggesting a potential link between SGSH activity and cell cycle control.

Since CDK11B and CDK5R1—also miR-4671-5p targets—are known cell cycle regulators, we evaluated whether miR-4671-5p overexpression influenced cell cycle progression. We observed a significant increase in S-phase cells and a reduction in G2-phase cells (**Figure 4G**), indicative of intra-S-phase arrest. While this is consistent with the known roles of CDK11B and CDK5R1 in S–G2 transition, SGSH may also contribute to this effect through HS accumiation. Heparan sulfate accumulation is known to inhibit topoisomerase I, which disrupts DNA replication and induces intra-S-phase arrest (Kovalszky et al., 1998; Lin et al., 2014).

Thus, miR-4671-5p may induce intra-S arrest through dual mechanisms: direct repression of cell cycle kinases and indirect effects mediated by SGSH inhibition and subsequent glycosaminoglycan accumulation. However, the precise contribution of SGSH to this arrest phenotype—and its interaction with canonical cell cycle regulators—warrants further investigation to clarify its mechanistic role in this regulatory axis.

In conclusion, our study identifies a novel Δ40p53–miR-4671-5p–SGSH regulatory axis that contributes to maintaining proper cell cycle progression. Under physiological conditions, Δ40p53 suppresses miR-4671-5p, allowing sustained expression of SGSH and other cell cycle regulators such as CDK11B and CDK5R1, thereby supporting controlled S-phase progression. However, in pathological conditions where Δ40p53 levels are reduced, miR-4671-5p is derepressed, leading to downregulation of its targets and activation of intra-S-phase arrest. These findings emphasize the importance of isoform-specific p53 signaling in regulating non-coding RNA networks and suggest that alterations in Δ40p53 expression could have broad implications for cancer biology and potential therapeutic targeting (**Figure 4H**).

## ACKNOWLEDGEMENTS

We thank Prof. J.C. Bourdon (University of Dundee) and Dr. Robin Fahraeus (INSERM) for the anti-p53 antibody. We acknowledge NIBMG (Kalyani, India) for RNA sequencing facilities and the SD lab members for helpful discussions. This work was supported by a DBT research grant to SD, who also received the J.C. Bose fellowship. Additional support came from the DBT-IISc partnership, DST-FIST Level II, and the UGC Centre of Advanced Studies.

## AUTHOR CONTRIBUTIONS

AP and SD: Conception and design of studies analysis, interpretation, and article writing. AP, SKT and PKG: performing experiments, interpretation of results and article editing. DK: bioinformatic analysis, interpretation of results, article writing and editing. SaG, SP and AM: RNA sequencing and analysis. SD: Funding acquisition, supervision and project management.

## DATA AVAILABILITY STATEMENT

The sequencing data files obtained is available in the online biorepository forum with the SRP BioProject ID **PRJEB47067**.

## FIGURE LEGENDS

**Figure S1:** (Table 1): miR-4671-5p targets obtained from TargetScan 7.2 database and miRDB. **(Table 2)**: Result of PANTHER Statistical Overrepresentation Test of “GO molecular function”-terms obtained from 786 predicted miR-4671-5p targets, using all *Homo sapiens* genes as reference list. FDR: False Discovery Rate. Results for FDR P < 0.05 reported. **(Table 3)**: Survival analysis of patients as classified by SGSH mRNA expression levels in pancreatic adenocarcinoma, colon tumor, glioblastoma and neuroblastoma gene expression datasets available on R2: Genomics Analysis and Visualization Platform. p-values <0.07 reported, below 0.05 considered significant. Similar analysis for hsa-miR-4671-5p expression, as available in the sole miRNA expression dataset on R2.

